# Maternal iron deficiency impacts the placental arterial network

**DOI:** 10.1101/2021.05.06.442902

**Authors:** Jacinta I. Kalisch-Smith, Emily C. Morris, Mary A.A. Strevens, Andia N. Redpath, Kostantinos Klaourakis, Dorota Szumska, Jennifer E. Outhwaite, Joaquim Miguel Vieira, Nicola Smart, Sarah De Val, Paul R. Riley, Duncan B. Sparrow

## Abstract

Placental vascular gene networks in mammals have been largely unexplored due to a lack of well validated molecular markers to identify them. This is required to study how they form in development, and how they are impacted by embryonic or maternal defects, which in-turn adversely affects the forming heart and vasculature. Such defects are known to be a consequence of maternal iron deficiency (ID), the most common nutrient deficiency world-wide. Here we employed marker analysis to characterise the arterial/arteriole and venous/venule endothelial cells (ECs) during normal placental development, and in the context of maternal ID. We reveal for the first time that placental ECs are unique compared with their embryonic counterparts. In the developing embryo, arterial ECs express *Neuropilin1* (*Nrp1*), *Delta-like ligand 4* (*Dll4*) and *Notch1*, while developing venous ECs express *Neuropilin2* (*Nrp2*), *Apj (Aplnr)* and *Ephrinb4 (Ephb4)*. However, in the E15.5 placenta, *Nrp1* and *Notch1* were restricted to arteries, but not continuing arteriole ECs. The arterial tree exclusively expressed *Dll4. Nrp2* showed pan-EC expression at E15.5, while *Ephb4* was not present at this stage. However, we found the placental venous vascular tree could be distinguished from the arterial tree by high versus low Endomucin (EMCN) and *Apj (Aplnr)* expression respectively. Using EMCN, we reveal that the placental arterial, but not venous, vascular tree is adversely impacted by maternal ID, with reduced area, total length and number of junctions of all vessels without affecting the EMCN high vessels. Defects to the embryonic cardiovascular system can therefore have a significant impact on blood flow delivery and expansion of the placental arterial tree.

## Introduction

The developing embryo requires oxygen, nutrients and disposal of waste products to facilitate its ongoing development. The placenta is the primary organ for facilitating these needs from mid-gestation in the mouse, but this takes place earlier in humans. In the mouse, the labyrinth vasculature provides an interface between mother and foetus. Situated on the embryonic side of the placenta, it is made up of arteries and veins which differentiate from the allantoic mesenchyme, originally residing in the embryo. Morphogenesis of this vasculature begins at embryonic day (E)7.5 whereupon several stem cell populations feed into the allantoic bud, (such as extraembryonic visceral endoderm and the primitive streak, and transforms into a branched endothelial cell (EC) network expressing PECAM1 (also known as CD31) (Drake, et al. 2000, Downs and Rodriguez, 2020). These ECs penetrate the base of the placental chorion, branch and form one layer of the interhaemal membrane (Simmons, et al. 2008, Cross, et al. 2003). The placental vascular trees have been identified by studies using plastic vascular casts, or microcomputed tomography imaging. The venous vascular tree is less complex than the arterial tree, containing 20% fewer vessels and smaller end-point capillaries than arterial trees at E15.5 (Rennie, et al. 2017). These vascular trees are also known to change dramatically in volume and branching complexity after environmental perturbations and genetic knockouts which impact blood flow from the heart (Detmar, et al. 2008, Bainbridge, et al. 2012, Withington, et al. 2006, Adamson, et al. 2002). However, we currently have no way to identify and define the cells and structural components of the placental vascular tree such as arterial ECs, venous ECs, and pericytes.

Analysis of the placental arterial and venous EC vasculature has stalled in recent years due to a lack of validated genetic markers to identify them. Little information exists of how they differentiate, the genetic programs they use prior to the onset of flow and during maturation, and whether they are molecularly distinct from other vascular beds in the embryo. Two subsets of ECs have been identified at E10.5. Candidate markers *Vegfa* and *Vegfc*, as well as the Apelin receptor *Apj (Aplnr)* have been proposed as arterial and venous markers, respectively. In the embryo, these are well defined markers of arterial and venous specification, and so may be also acting in a similar manner in the placenta. The placental endothelium may also express other genes similar to the systemic embryonic endothelium, such as vein EC-specific Eph receptor B4 (*EphB4)* and artery EC-specific Ephrin B2 (*Efnb2)*. These are expressed at E9.5 in the umbilical vein and artery respectively (Wang, et al. 1998), which connect to placental arterial and venous vessels, and are also allantoic-derived. Little is known about venous differentiation in the placenta. Recent single cell analysis of labyrinthine cells showed unique cell type clustering of allantoic mesenchymal and ECs, and an additional cluster marked by *Podoplanin* (*PDPN*) expression (Marsh and Blelloch, 2020). However, this study did not distinguish between arterial and venous ECs, nor had it elaborated on the nature of the PDPN enriched cluster.

Here, we describe distinct marker expression in 3 individual labyrinthine cell types: arterial ECs, venous ECs and pericytes using protein and RNA localisation. We then apply these markers to a model of maternal iron deficiency (Kalisch-Smith, et al. 2019, pre-print) which results in impairments of both the embryonic cardiac and lymphatic system. In embryos presenting with these cardiovascular defects, matched placentas have reduced growth of the arterial EC tree but not the venous EC tree. We also show common lymphatic genes are used by labyrinthine blood vessels.

## Results

### Investigation of embryonic arterial and venous gene expression in the placenta

We first profiled common embryonic markers for arterial ECs, venous ECs and pan-EC markers (Chong, et al. 2011) in the placenta. We focused on different types of structures which transport blood through the placenta, i.e. from the umbilical circulation (systemic artery/vein), through to the arterial stems (primary branches of vascular tree) and to terminal branches of capillaries (arterioles and venules). The E15.5 timepoint was used because the placenta at this stage has undergone the majority of growth and differentiation. Expression profiles were compared to either CD31/PECAM and Isolectin B4 (ILB4), which are pan-EC markers.

Firstly, we investigated NRP1 and NRP2. In the embryo, these are known to be specific to arterial and venous ECs respectively (Chong, et al. 2011). However, the placental vasculature showed a different expression pattern. NRP2 was present in a subset of CD31 positive ECs throughout the placenta at E15.5, while NRP1 was found in larger vessels (most likely arterial based on their size) at E13.5 (Fig.1A,B), but not E15.5 (data not shown). Both NRP1/2 were expressed in the allantoic mesenchyme (Fig.1A,B). In the embryo, endomucin (EMCN) is restricted to capillaries and venous ECs from E15.5 in development (Brachtendorf et al., 2001). In the placenta, EMCN showed distinct expression patterns to pan-EC ILB4 and CD31 at E12.5 and E15.5 respectively (Fig.1C, Fig.S2G). At both time points, high expression can be seen in collecting vessel ECs at the base of the labyrinth (Fig.S2G). Later, at E15.5, EMCN showed higher expression in venules within the lower portion of the labyrinth, compared with arterioles within the upper portion. Delta-like ligand 4 (*Dll4)*, a Notch ligand, is a robust embryonic arterial EC marker from E8.0 (Chong, et al. 2011), and is also present in the umbilical and vitelline arterial ECs and some placental vessel ECs at E9.5 (Duarte, et al. 2004). In addition, *Dll4* null mice show embryonic lethality by E10.5, with aberrant vessel formation and regression of the placental vasculature (Arora, et al. 2012, Duarte, et al. 2004, Gale et al. 2004). In the placenta, *Dll4* mRNA was more abundantly expressed in the umbilical arterial ECs than the vein ECs (Fig.1D, D’’’ vs D’’’’), and detected in arteriole capillaries (see Fig.1D, D’, D’’), colocalising with *Cd31* mRNA. This agrees with the embryonic expression outlined above. Another interesting expression pattern was shown by DACH1. DACH1 is a transcription factor that has been previously shown to promote EC migration and coronary artery growth (Chang et al. 2017), and more recently, implicated in pre-artery specification and differentiation of arterial ECs (Raftrey and Red-Horse, 2020, pre-print). In the placenta at E15.5, DACH1 showed strong expression in the ECs of collecting vessels (Fig.1E,E’’, Fig.S2G) and nuclear expression in capillaries extending towards the junctional zone (Fig.1E’). *Apelin receptor (Aplnr/ Apj)* is expressed at E9.5, restricted to one vessel leading to the placenta, and is known to facilitate vessel sprouting and branching (Freyer, et al. 2017). Curiously, APJ is a venous marker in the retina (Saint-Geniez, et al. 2002, cardinal vein (Chong, et al. 2011), and the sinus venosus, a progenitor to the coronary vessels (Sharma, et al. 2003). *Apj* is specific to allantoic endothelial cells and it can signal to *Ela* on the adjacent trophoblast in a paracrine manner (Ho, et al. 2017). *Apj* null mice exhibits vascular defects in the embryo and embryonic death from E10.5 (Kang, et al. 2013). *Apj* mRNA puncta were highly localised to the umbilical vein ECs but not to the umbilical artery ECs or central arterial stems (Fig.1F). Similar to the patterns of EMCN expression, *Apj* was more highly expressed in basal venous ECs, but was also present in arteriole ECs. See Fig.1G for diagrammatic summary of expression profiles. Negative control staining for mRNA showed background staining for the Cy3 channel, and no staining for Cy5 (Fig.S2H). Taken together, this analysis shows that the placenta only mimics a subset of embryonic expression patterns, and is a unique endothelial organ bed.

**Figure 1.**
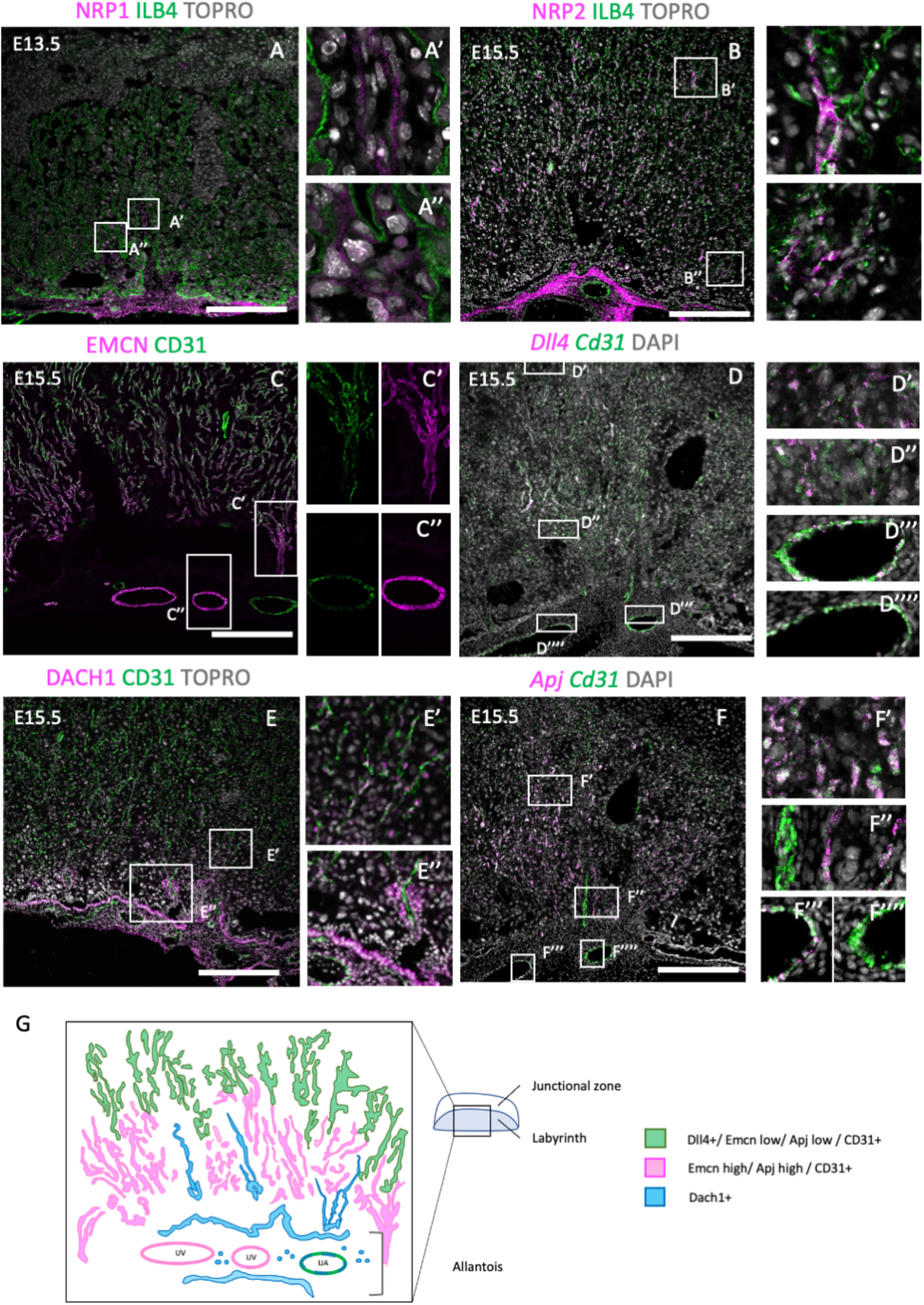
Investigation of embryonic arterial and venous genes during mouse placentation. E13.5 or E15.5 placentae were stained with embryonic arterial-specific genes (NRP, DACH1, *Dll4*, A, D, E), and venous-specific genes (NRP2, Endomucin (EMCN), *Apj* in B, C, F). The boxes outline the areas shown at higher magnification in in A-F (‘). (G) Diagrammatic representation of arterioles (green), arterial stems and allantoic mesenchyme (blue) and venous (magenta) genes within the E15.5 placenta. Scale bars (white) = 200um.

In order to further understand labyrinth morphogenesis, we conducted further analysis of key vascular genes in the placenta at earlier developmental time points. CD105 (Endoglin) showed strong pan-EC expression at E12.5, consistent with its role in proliferating endothelial cells. (Fig.S2A). Alpha smooth muscle actin (αSMA) was found in labyrinthine and allantoic mesenchymal cells at E10.5 (Fig.S2B), and in a central artery projecting to the upper portion of the labyrinth. This expression pattern remained the same at E12.5 (Fig.S2C). *Notch1* expression has been previously identified in the embryo in arterial ECs and shows expression by E8.25 (Chong et al. 2011). In the placenta, NOTCH1 expression was primarily expressed in the smooth muscle of the umbilical artery at E15.5 (Fig.S2D), while SM-MHC (smooth muscle) was expressed highly throughout the allantoic mesenchyme, collecting vessels, and surrounding arterioles (Fig.S2E).

### Endothelial enhancer expression in the placenta

We next traced arterial and venous vessels back to earlier stages of placental formation. E10.5 in the placenta follows attachment of the trophoblast-derived chorion with the allantois, and initial branching to create a primitive vascular network. To investigate arterial and venous formation of the placenta, a 3D model was first created of the E10.5 placenta by high resolution episcopic microscopy (HREM) and volume reconstruction. Tracing of allantois-derived vascular structures showed two separate vascular trees emerging from the allantois and branching towards the chorion (Fig.2A-C). Vessels were segmented easily into upper (arterial, blue, deoxygenated) and venous (lower, red, oxygenated) portions. The E10.5 labyrinth was next assessed for sub-populations of allantoic-derived EC. We used well described and unique embryonic EC enhancer:LacZ constructs (Sacilotto, et al. 2013, Neal, et al. 2019) to identify if they are present in placental ECs. This is also the necessary first step to investigate each of their upstream regulatory pathways. The enhancer *Dll4-12:LacZ* is specifically active in arterial ECs, while *Dll4in3:LacZ* is expressed by both arterial and angiogenic ECs. Beta-Galactosidase staining showed positive localisation for both of these enhancers in ECs of the E10.5 placenta (Fig.2D-F). These cells formed small vascular structures bordering the labyrinth trophoblast, extending from the allantois at branching points. Both enhancers also showed some expression in the allantoic mesenchyme. In the embryo, the Ephb4-2:LacZ transgene is selectively expressed in venous ECs (Neal, et al. 2019). However, in the placenta it is active in the allantois and allantoic mesenchyme entering the labyrinth (Fig.2E). A subset of these cells was ILB4 positive, suggesting they could be a progenitor population. See Fig2G for summary of marker expression.

**Figure 2.**
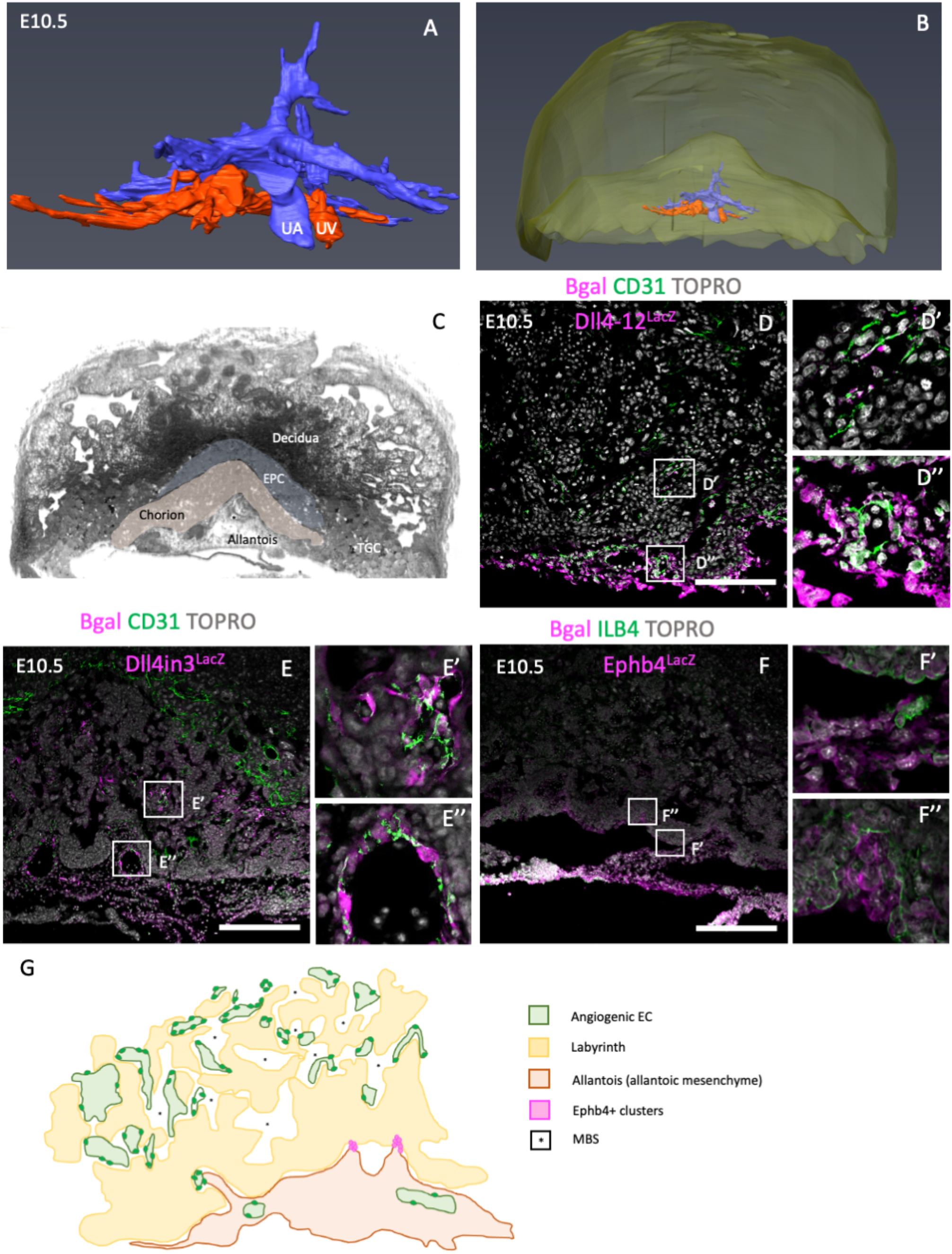
Vascular marker analysis of mouse placental labyrinthine cell types at E10.5. (A,B) 3D ultrastructural model of the arterial (blue, deoxygenated) and venous (red, oxygenated) vasculature of the E10.5 placenta, with Decidua (yellow) in (B). (C) shows midline section captured from high resolution episcopic microscopy (HREM), with pseudo-colouring for placental zones. (D) Bgal expression of enhancer *Dll4-12:LacZ*. (E) Bgal expression of *Dll4in3:LacZ*. (F) Bgal expression of *Ephb4:LacZ*. The boxes outline the areas shown at higher magnification in D-F (‘). (G) Diagrammatic representation of E10.5 placenta expression patterns for Bgal/LacZ enhancers. Bgal; B-galactosidase. Scale bars (white) = 200um. TGC; trophoblast giant cells.

### Common lymphatic genes are expressed in the placenta

The mucin-type protein podoplanin (PDPN) has been previously associated with epithelial-mesenchymal transition (EMT), and is expressed in lymphatic endothelium and epicardial cells (Thiery, 2002). In the placenta, PDPN was expressed in basal labyrinth cells (ILB4-, Fig.3A) at E10.5, with continuing cells leading into the midline becoming ILB4+ ECs (Fig.S2B). Similar expression patterns were shown at E12.5 (Fig.S2A). By E14.5/E15.5 a subset of these PDPN+ cells were also ILB4+, and were seen in major vessels extending towards the junctional zone, small crypts near the base of the labyrinth, and in allantoic mesenchyme surrounding embryonic blood vessels (Fig.3B,C, C’’’). Co-staining showed PDPN+ cells were adjacent to both EMCN+ blood vessels and NG2+ pericytes (Fig.3D,E). These placental PDPN+ cells could be therefore a subset of mesenchymal and blood vessels undergoing EMT, as they are dispersed throughout the labyrinth. See diagrammatic representations for expression patterns at E10.5 (Fig.3F) and E14.5 (Fig.3G).

**Figure 3.**
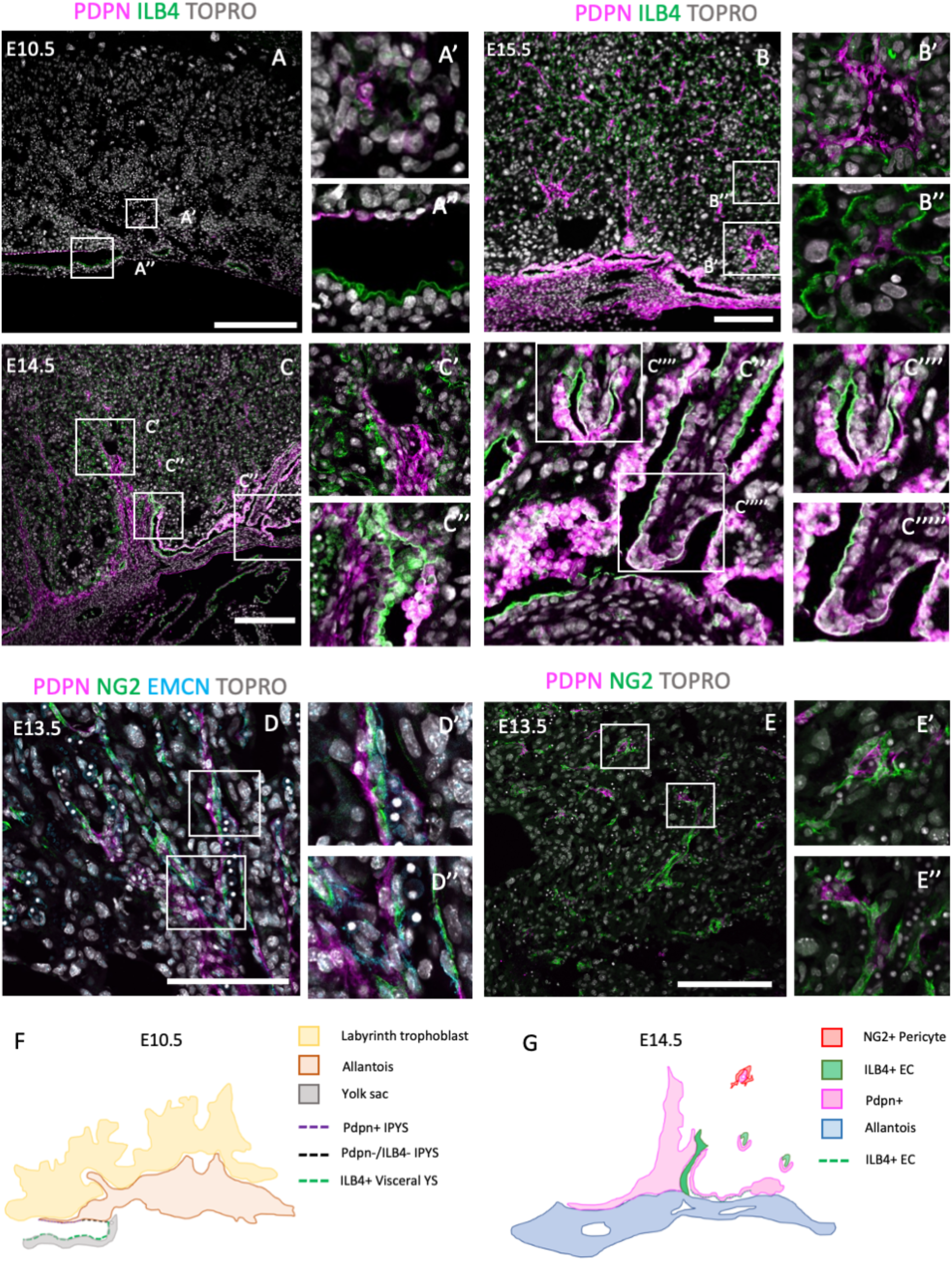
Investigation of common embryonic lymphatic genes during placentation. Podoplanin (PDPN, A-E) was investigated from E10.5 to E15.5 in combination with endothelial markers ILB4 and CD31, venous marker (EMCN high, arterial EMCN low), and pericyte marker NG2 (chondroitin sulphate proteoglycan). The boxes outline the areas shown at higher magnification in A-E (‘). Diagrammatic representations of PDPN expression at E10.5 (F) and E14.5 (G). Scale bars (white) = 200um.

Lymphatic vessel endothelial hyaluronan receptor-1 (LYVE1) has been previously reported to be expressed in embryonic blood vessels, lymphatic vessels and the lymph node, among other organs (Gordon, et al. 2008). *Lyve1* is reported to be expressed in labyrinthine ECs, albeit only at E11.5/E12.5 (Shaut, et al. 2008). To investigate this, we used the *Lyve1-Cre:TdTomato* system (Pham, et al. 2009). TdTomato+ cells were localised in visceral yolk sac and allantoic mesenchyme at E10.5, and in all labyrinthine ECs to E14.5 (Fig.S3A, B, D, F). Yolk sac hemogenic endothelia have been previously reported as *Lyve1+* (Lee, et al. 2015). The TdTomato system has been known to over-report expression in highly expressed genes, so we therefore decided to compare our reporter with antibody/protein localisation. Antibody staining partially matched the *Lyve1* reporter, indicating blood vessel ECs at E13.5 (Fig.S3C), but were absent in ECs at E10.5 and E15.5 (Fig.S3A,E). Antibody staining at E15.5 showed sporadic cells in the labyrinth and junctional zone, most likely to be macrophages. No expression of PROX1 was found (data not shown). Now that we have described *Dll4* marking placental arterial ECs, and EMCN/Apj as marking venous ECs, *Lyve1+* is therefore a pan-ECs marker in the placenta.

### Maternal iron deficiency impacts placental formation

Having defined markers of placental arterial and venous ECs, we wanted to apply them to a disease model to see how impairing the embryonic cardiovascular system can affect their formation. This is important as perturbation of either the placenta or the heart can impact the other, as shown in mouse mutant models (Perez-Garcia et al. 2018) and human epidemiological studies (Araujo Junior, et al. 2016). The heart must beat against the resistance of the placental vascular bed (Thornburg, et al. 2010), and therefore changes to placental blood flow are likely to impact development of the embryonic heart. As the placenta and heart develop in parallel, it is important to determine whether the heart defects are primary or secondary to placental defects. We have recently shown that maternal ID causes congenital heart defects and sub-cutaneous oedema (see Fig.S1). Considering these cardiovascular defects, it is unknown how they may impact placental growth, or vice versa.

We first assessed placental tissues for weight from E9.5 to E15.5 (Fig.4A). Placental weights were reduced between E9.5 and E15.5 as a result of ID. This was most significant at E12.5 (p<0.0001), coincident with the beginning of embryonic death in approximately half of embryos. The cause of this lethality is yet to be fully explored, and therefore could be due to placental insufficiency. Placental weights of surviving embryos remained reduced at E14.5 and E15.5 (p<0.01), albeit to a lesser extent. This is likely due to survivor effects of the remaining embryos whereby nutrients are reabsorbed from lethal embryos. Gross placental volumes were assessed at E9.5, E12.5 and E15.5. E9.5 placental tissue from somite-matched conceptuses was assessed by HREM (Fig.S4), and quantified for total volume and placental compartments; decidua, parietal trophoblast giant cells (P-TGCs), ectoplacental cone (EPC), chorion, and allantois. No significant changes were found in any placental compartment. We further assessed the E12.5 placenta after ID and found that while the whole placental trended towards a decrease in volume (p=0.0609, Fig.4C), the labyrinth vascular compartment was significantly reduced (p<0.001, Fig.4D). The junctional zone and decidua remained unchanged (Fig.4E,F, see Fig.4B for diagrammatic representation of placental zones). By E15.5, quantification showed ID reduced volumes of the whole placenta (p=0.011, Fig.4G), labyrinth (p=0.0057, Fig.4H), and decidua (p=0.0057, Fig.4J). No change was found in the junctional zone compartment (Fig.4I). Embryonic capillaries (blood spaces) were identified using the marker ILB4 (Fig.4L,M). Maternal blood space was markedly reduced after ID (p<0.0001, Fig.4K). Maternal blood spaces were identified by an absence of staining, and the presence of large nuclei of sinusoidal trophoblast giant cells lining these blood pools.

**Figure 4.**
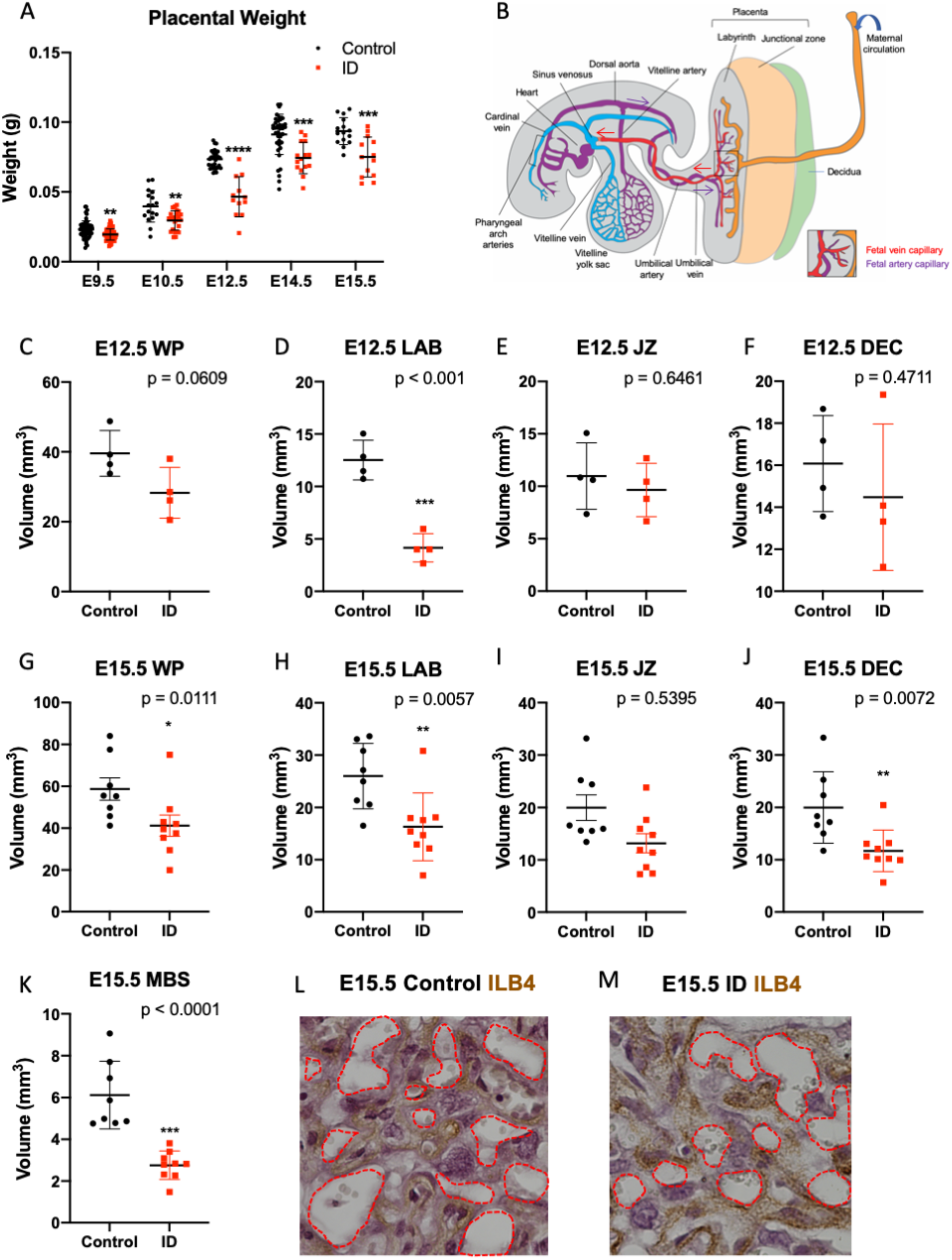
Maternal iron deficiency impacts placental formation, particularly to the labyrinth vasculature. (A) Placental wet weights from E9.5 to E15.5. After ID, placentas weighed less, most significantly at E12.5 (onset of embryonic death). Placental weights remained reduced to E15.5. (B) Diagrammatic representation of placenta-foetal blood flow in the mouse in mid-gestation with maternal blood spaces (orange), and foetal capillaries made up of PA (purple) and PV (red). Arrows show direction of blood flow. (C-F) Gross morphological volumes of placental compartments at E12.5 for whole placenta (WP, C), labyrinth (LAB, D), junctional zone (JZ, E) and decidua (DEC, F). Labyrinth volume only was reduced after ID. (G-L) Gross morphological volumes of placental compartments at E15.5. Whole placental volume was reduced (G), including compartments for the labyrinth (H) and decidua (J). Maternal blood spaces (MBS, K) were quantified as Isolectin B4 (ILB4) negative staining, and was reduced at E15.5. ILB4 staining of E15.5 placentas for control (L) and ID (M), with red dashed line demarcating MBS. (A) Control: E9.5 n=58, E10.5 n=17, E12.5 n= 32, E14.5 n=40, E15.5 n=15. ID: E9.5 n=59, E10.5 n=24, E12.5 n=12, E14.5 n=15, E15.5 n=13. (C-F) Control: n=4, ID n=4. (G-K) Control: n=8, ID: n=9. Control (black dots), ID (red squares). Data represented as mean ± SD. Data was analysed by unpaired t-test unless otherwise stated. Mann-Whitney test was performed in A (E10.5 only), G-I. *P<0.05, **P<0.01, ***P<0.001, ****P<0.0001. ID; Iron Deficient.

### Maternal ID impacts placental arterial but not venous networks

We next sought to assess placental arterial and venous vasculature after ID, stratifying for those from embryos with and without heart defects and oedema (H+O), see Table.S1, Fig.S5 for summary of heart defects, HREM image stacks for representative control; Movie1 or ID; Movie2. Despite these defects, ID did not impact ventricular volume (Fig.S5a). Midline sections of placentae were stained for EMCN and counted for EMCN high (venous ECs) and total ECs (CD31+). ID caused a reduction in the total area, number of junctions, and total length of CD31+ vessels (Fig.5A-C, see representative images for control Fig.5K, and ID placentas Fig.5L). Post-hoc analysis revealed this effect in Fig.5B and C was driven by ID H+O embryos (p<0.05 vs control), and not by ID embryos with none. ID H+O placentas also showed greater variation than ID embryos with no defects. PDPN+ blood vessels were also assessed but were unchanged with area, junctions or length (Fig.5D-F). Finally, EMCN high venous vessels were assessed for area, junctions and total length after ID or ID H+O, but found no differences (Fig.5G-I). Surprisingly, ID embryos with no defects increased the total length of EMCN-high vessels compared with controls (Fig.5I). Since ID reduced total EC vessels without changing venous vessels, this suggests that ID impacts primarily growth of the placental arterial tree. The placental arteries receive high pressure blood pumped from the heart to the remaining embryo. Therefore, heart defects causing reduced blood flow to placenta are likely to affect arterial expansion. However, given that the placental weight was markedly reduced at E9.5, prior to any physiological changes in the heart, this suggests that the placental defect could be initiating later embryonic cardiovascular defects.

**Figure 5.**
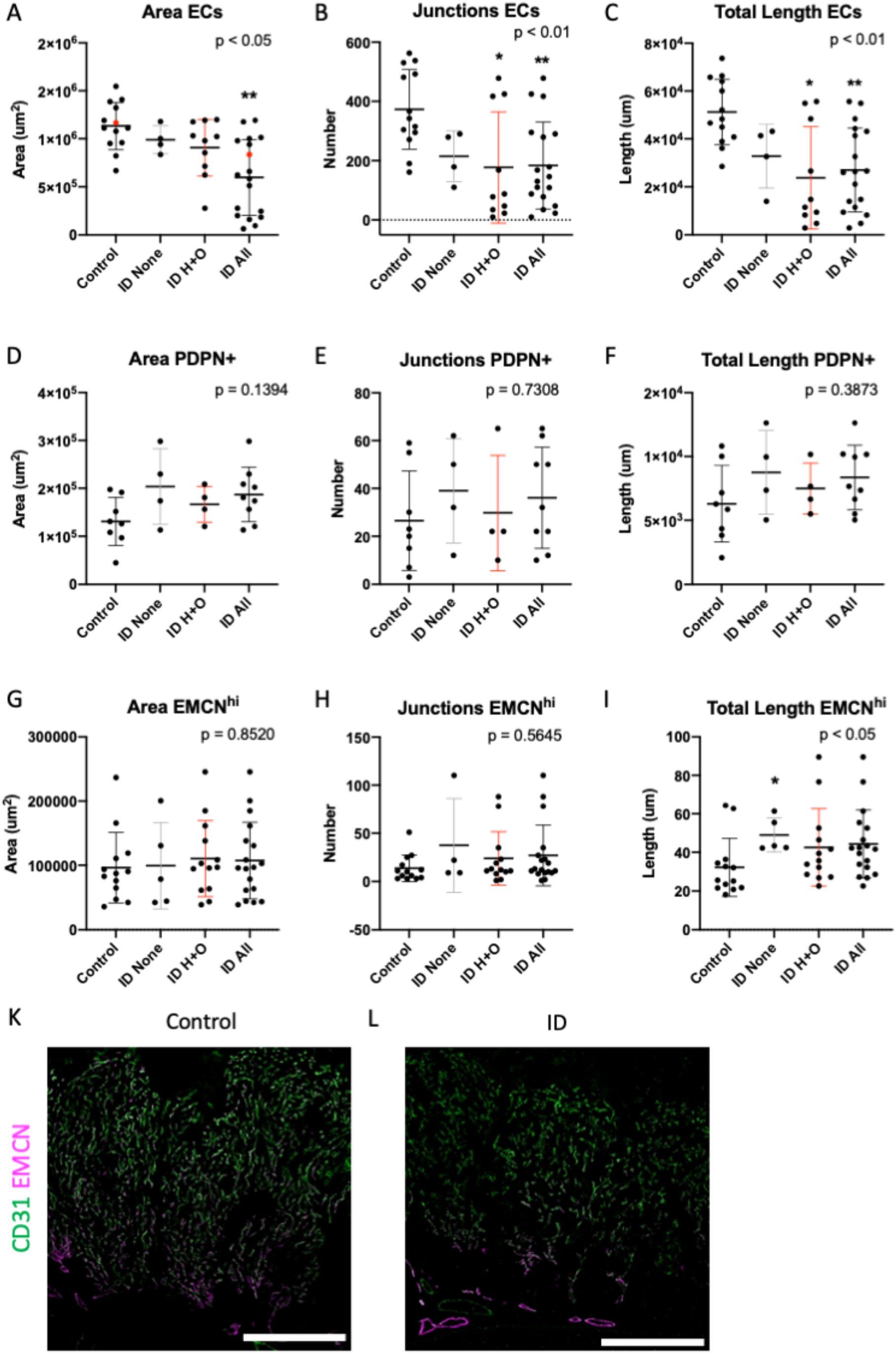
Maternal iron deficiency reduces total but not venous-specific ECs in the E15.5 placenta. Total ECs (A-C), PDPN+ ECs (D-F) and veins (EMCN^hi^) (G-I) were assessed for vessel area, number of junctions and total length. ID samples were stratified for those with heart defects and oedema (ID H+O) or those with none (ID none). Red points in (A) correspond to placental images in (K, control) and (L, ID), showing CD31 (pan-EC) and thresholded EMCN to highlight strong venous expression. Data represented as mean ± SD. Data was analysed by ordinary one-way ANOVA if normally distributed (D-F), or if not by Kruskal-Wallis test (A-C,G-I). Post-hoc analysis was compared to the control group *p<0.05, **P<0.01. Control n= 13 (A-C,G-I), 8 (D-F). ID n=18 (A-C,G-I), 9 (D-F). ID; Iron Deficient. Scale bars (white) = 200um.

## Discussion

Here, we describe placental arterial and venous EC genetic marker profiles, which are unique compared with the embryo. Curiously, umbilical/placental arterial stems and venous profiles differed to their labyrinthine capillary derivatives; arterioles and venules. We then used these newly established profiles to study a mouse model of maternal iron deficiency (ID), which resulted in cardiovascular defects and oedema (Kalisch-Smith et al. 2020, pre-print). Curiously, in the present study, placental analysis showed reduced labyrinthine development only at E12.5. Models restricting blood flow to the placenta that cause perturbed labyrinth formation are often associated with heart defects. These include knockout of *Ncx1*, a cardiac Na^+^/Ca^2+^ exchanger, which impacts embryonic heart rate (Cho et al. 2003), and therefore blood flow. In addition, administration of excess glucocorticoids has also shown reduced placental vascularisation that coincides with delayed heart development (Wyroll, et al. 2016). Many genetic knockout models show heart and placental defects such *Cxadr* (Outhwaite, et al. 2019), *Fltr2* (Tai-Nagara, et al. 2017), *Hoxa13* (Shaut, et al. 2008), p38a MAPK (Adams, et al. 2000), among others (reviewed by Maslen, 2018, Camm, et al. 2018).

In addition, we also found the placental arterial tree was markedly reduced at E15.5, driven by the embryonic cardiovascular defects. Defects specific to the placental arterial tree are commonplace. Other models including hypomorphic *Gcm1* expression (Bainbridge, et al. 2012) and exposure of mouse dams to cigarette smoke (polycyclic aromatic hydrocarbons) (Detmar, et al. 2008), both increased the placental arterial tree. Knockout of *Cited2*, however, effects both arterial and venous trees (Withington, *et al*. 2006). Previous investigation by Navankasattusas, et al. (2008) showed that a reduction in placental arteriole formation was associated with blood flow reversal to the embryo, and was the cause of lethality at E13. This could similarly occurring in our ID model. It is currently unknown what teratogens, if any, might impact primarily the venous placental vasculature.

Evidence that a link between congenital heart defects (CHD) and placental perturbations in humans is emerging. In a study of 924,422 liveborn Danish babies, they showed that CHD was associated with smaller placental size at birth, teratology of Fallot, DORV, and VSD (Matthiesen, et al. 2016). Additional CHDs have been associated with placental vascular defects, including hypoplastic left heart syndrome (Jones et al., 2015, Rychik et al., 2018, Courtney, et al. 2020). Maternal iron deficiency is the most common nutrient deficiency of women of reproductive age, and affects approximately 500 million women world-wide (WHO, 2015). It is therefore possible that iron deficiency is contributing to human incidences of CHD. Our previous investigation into CHD showed that iron supplementation can be used to rescue the observed heart defects (Kalisch-Smith, et al. 2020, pre-print). Since the heart defects in our model impact placental formation at least in part, this research would advise supplementation of iron to women planning and during pregnancy to prevent heart-placental defects.

## Conclusion

This study showed that the placenta is a unique endothelial bed expressing only a subset of embryonic EC markers. While true placental venous markers remain elusive, we have shown that venous ECs/venules express higher EMCN and *Apj* than their arterial counterparts. These markers can be used to better understand placental development and dysfunction. We have investigated one common maternal condition; maternal iron deficiency, and shown an arterial-specific defect in the placenta. Further research is required to understand how embryonic defects impact the placental vasculature, particularly to venous formation.

## Acknowledgements

This research was supported by a BHF Senior Basic Science Research Fellowship FS/17/55/33100 (DBS); Oxford BHF CRE core infrastructure grants (DBS and 1225 JIKS, RE/18/3/34214).

## Author contributions

Conceptualisation; JKS, DS. Methodology; JKS, DS, AR. Formal analysis; JKS, ECM, MAAS, ANR, JO, DSz. Investigation; JKS, ECM, DSz, KK, AR, JV. Resources; DS, NS, SdV, JV, PR. Writing original draft; JKS. Reviewing and editing; JKS, JV, NS, DS, ANR, PRR, MAAS, DSz, SdV. Visualisation; JKS, ECM, MAAS. Supervision; JKS and DS. Project administration; JKS, DS. Funding acquisition; DS, JKS.

## Methods

### Animal housing and diet modification

All animal experiments were compliant with the UK Animals (Scientific Procedures) Act 7391986, approved by the University of Oxford animal welfare review board and the Home Office (project license PB01E1FB3). Mice were housed in an SPF facility with a 12:12 hour light/dark cycle, at 19-23°C, 55 ±10 % humidity, with free access to food and water. Bedding was changed fortnightly, and animals were assessed daily for welfare. Mice were fed with standard chow TD.08713 (control), or TD.99397 (iron deficient) feed (Envigo, Belton, UK). C57BL/6J mice were purchased from Charles River UK. The following genetically-modified mouse strains were used; *Lyve1*-Cre (JAX, 012601, *Lyve1*^*tm1*.*1(EGFP/cre)Cys*^), R26R-tdTomato (Madisen, 2010). Enhancer mouse lines used as previously described; Tg(Dll4in3:*lacZ*) (Neal, 2019), Tg(Dll4-12:*lacZ*) (Sacilotto, 2013), EphB4-2:*lacZ* (Neal, 2019).

### HREM (High Resolution Episcopic Microscopy) assessing heart and placental morphology

Somite-matched placentas at E9.5 were fixed in 4% paraformaldehyde and processed in a methanol series (10%, 20%, 30%, 40%, 50%, 60%, 70%, 80%, 90%, 95%, 100%) for 1-hour washes each. Samples were incubated in 50:50 mixture of methanol and JB4 resin (Polysciences, 00226-1, GMBH, Germany) overnight. Samples were incubated in JB4 resin for 1h, and transferred to fresh resin for incubation overnight. E15.5 embryos were fixed overnight in 4% paraformaldehyde at 4°C, and were dissected for heart-lung complexes. These samples used the same conditions as above, except the final JB4 incubation, which was for 3 days. Samples were embedded individually in JB4 according to manufacturer’s instructions. Samples were cut (2um E9.5 placenta, 3um E15.5 heart-lung complex) on an optical HREM (OHREM) microscope (Indigo scientific) with images taken using a Jenoptik Gryphax camera. Image stacks were processed into cubic data and reduced to either 10% or 50% for 3D modelling using the Amira software package 2019.4 (Thermo Fisher Scientific). Greyscale images were imported into MD DICOM viewer version 9.0.2 (Pixmeo), or Horos 3.3.6 (https://horosproject.org) in a 50% stack, inverted to black-white, and rendered into 3D.

### Immunohistochemistry (IHC), imaging and placental quantification

IHC on paraffin sections was performed as previously described (Shi, et al. 2016). Placental tissue was fixed in 4% paraformaldehyde overnight, bisected with a double-edged blade, with half for paraffin embedding and half for cryosectioning, both embedded cut face down. Paraffin embedded tissue was cut at 7um. ID samples at E12.5 and E15.5 were used for stereology as described by Coan et al. (2004). E12.5 tissue was sampled at 1:25, while E15.5 tissue was sampled at 1:50 sections. Slides were dewaxed with xylene and rehydrated with an ethanol series prior to antigen retrieval in either TE pH8.5 or citrate buffer for 20 minutes, with 20-minute cool down. Slides were processed using a Shandon Sequenza® Immunostaining Center (Thermo 827Fisher Scientific). Slides were blocked in 2.5% donkey/goat serum in PBST for 45 minutes at RT, then in Sudan black for 20 minutes to removed autofluorescence. Slides were washed in 3x PBST prior to incubation in primary antibody (in 2.5% block in PBST) overnight at 4°C. Slides were washed 1x PBST prior to incubation in secondary antibody (in 2.5% block in PBST) for 1h. For fluorescent slides, this included TO-PRO®-3 Iodide nuclear dye (T3605, Life Technologies 1:10,000). Slides were washed 3x PBST prior to mounting with anti-fading PVA/Moviol-DABCO (Sigma) medium. Fluorescent slides were imaged using an Olympus FV3000 confocal microscope using an Olympus UPLSAPO NA0.4 10x objective. Individual images were captured at 1024×1024 pixel resolution.

For isolectin B4 staining (ILB4), slides underwent TE antigen retrieval, washed in PBS for 5 minutes, and blocked for endogenous peroxidases using 0.9% H2O2 in MQ H2O for 10 min. Slides were then incubated in 0.1% triton with 0.1mM ions (MgCl2, CaCl2, MnCl2) for 10 min prior to incubation with IsolectinB4 (derived from *Bandeiraea simplicifolia*) in PBS for 2h at RT. Slides placed in DAB substrate (Vector peroxidase substrate kit, SK-4100) for colour reaction. This reaction was stopped by washing in milliQ water. Following, slides were counterstained with haematoxylin and mounted in DPX medium. For light microscopy, whole placental sections were imaged at x1.25(obj) magnification using an Olympus SZX7 Compact Stereo Microscope (E12.5 & E15.5) and three fields of view of the labyrinth (E15.5) were imaged using a Nikon Eclipse Ci microscope at x40(obj) magnification. Placental compartments were then estimated using the Cavalieri principle, as described by Coan et al. (2004) and analysed in ImageJ (NIH). A 100 μm^2^ and 200 μm^2^ grid was respectively superimposed onto each field of the ⨯1.25 and ⨯40 sections and the number of points falling onto each placental compartment was counted, blind to treatment group. At E12.5, total, decidua, junctional zone and labyrinth were estimated. At E15.5, fetal and maternal blood spaces were estimated in addition to these compartments.

Counting of total and venous (EMCN^hi^) ECs and associated vessels was performed using Angiotools (Zudaire, et al. 2011). Image stacks (n=4) created an average intensity projection. CD31 images were used for total counts, while EMCN images were thresholded and cropped to only show venous positive vessels.

| Antibody | Species/<br>Conjugation | Clonality | Dilution | Company | Product Code | Antigen retrieval/<br>Cryosections |
| --- | --- | --- | --- | --- | --- | --- |
| a-SMA | Rabbit | Polyclonal | 1:200 | Abcam | ab5694 | CRYO |
| BGAL | Chicken | Polyclonal | 1:200 | Abcam | ab9361 | CRYO |
| CD105/<br>ENDOGLIN | Goat | Polyclonal | 1:25 | R&D Systems | AF1320 | CRYO |
| CD31 | Armenian<br>Hamster | Monoclonal | 1:100 | Abcam | ab119341 | CRYO |
| ENDOMUCIN | Rat | Monoclonal | 1:50 | Santa Cruz | sc53941 | CRYO |
| GATA4 | Mouse | Monoclonal | 1:100 | Santa Cruz | sc-25310 | CRYO |
| Isolectin B4 | Biotin | - | 1:200 | Sigma | L2140 | TE |
| Isolectin B4 | Peroxidase<br>conjugate | - | 1:100 | Sigma | L5391 | TE or Citrate |
| LYVE1 | Rabbit | Polyclonal | 1:100 | Angbio | 11-034 | CRYO |
| NG2 | Rabbit | Polyclonal | 1:100 | Millipore | AB5320 | Citrate |
| NOTCH1 | Rabbit | Polyclonal | 1:100 | Abcam | Ab8935 | CRYO |
| NRP1 | Rabbit | Monoclonal | 1:50 | Abcam | ab81321 | - |
| NRP2 | Rabbit | Monoclonal | 1:50 | Cell Signaling<br>Tech | D39A5 | CRYO |
| PODOPLANIN | Hamster | Monoclonal | 1:100 | Fitzgerald | 10R-P155a | TE, Citrate or<br>CRYO |
| SM-MHC | Rabbit | Polyclonal | 1:100 | Abcam | Ab125884 | CRYO |

### RNAscope

RNAscope Multiplex Fluorescent v.2 assay (ACD) was performed on 12um cryosections according to the manufacturer’s instructions with minor modifications (previously described by Lupu, et al. 2020). Slides were boiled for 8mins. Protease Plus digestion was performed at 30mins at RT. Probes were optimized for hybridization at 40°C. TSA plus fluorophores (Akoya Biosciences) used: Cy3 (1:1500), Cy5 (1:1500). Slides were counterstained with DAPI. The following catalogue probes were used: *Pecam1* (#316721-C3), *Aplnr* (*Apj*, #436171), *Dll4* (319971). The 3-plex negative probe against dapB was used as a negative control (#320878). Assays were run in duplicate. Confocal imaging was performed as outlined above, except 4x Z stacks were taken. Average intensity projections were used as representative images.

### Statistical analyses

All statistical analyses were performed with Prism 8.4.2 (GraphPad Software). Data were tested for normal distribution by Shapiro-Wilk tests. For data with two groups, normally distributed samples were analysed by two-tailed Student’s t test. Non-normally distributed samples were tested using a two-tailed Mann-Whitney test. For >2 groups, a parametric one-way ANOVA or non-parametric Kruskal Wallis test was used. Statistical significance was set at <0.05. Data are presented as mean ± standard deviation (SD).

### Data availability

HREM datasets are available upon reasonable request.

## References

Adams, R.H., Porras, A., Alonso, G., Jones, M., Vintersten, K., Panelli, S., Valladares, A., Perez, L., Klein, R. and Nebreda, A.R. (2000). Essential Role of p38α MAP Kinase in Placental but Not Embryonic Cardiovascular Development. Molecular Cell, 6(1), pp.109– 116.

Araujo Júnior, E., Tonni, G., Chung, M., Ruano, R. and Martins, W.P. (2016). Perinatal outcomes and intrauterine complications following fetal intervention for congenital heart disease: systematic review and meta-analysis of observational studies. Ultrasound in Obstetrics & Gynecology, 48(4), pp.426–433.

Arora, R. and Papaioannou, V.E. (2012). The murine allantois: a model system for the study of blood vessel formation. Blood, 120(13), pp.2562–2572.

Bainbridge, S.A., Minhas, A., Whiteley, K.J., Qu, D., Sled, J.G., Kingdom, J.C.P. and Adamson, S.L. (2012). Effects of Reduced Gcm1 Expression on Trophoblast Morphology, Fetoplacental Vascularity, and Pregnancy Outcomes in Mice. Hypertension, 59(3), pp.732–739.

Brachtendorf, G., Kuhn, A., Samulowitz, U., Knorr, R., Gustafsson, E., Potocnik, A.J., Fässler, R. and Vestweber, D. (2001). Early expression of endomucin on endothelium of the mouse embryo and on putative hematopoietic clusters in the dorsal aorta. Developmental Dynamics, 222(3), pp.410–419.

Camm, E.J., Botting, K.J. and Sferruzzi-Perri, A.N. (2018). Near to One’s Heart: The Intimate Relationship Between the Placenta and Fetal Heart. Frontiers in Physiology, 9.

Chang, A.H., Raftrey, B.C., D’Amato, G., Surya, V.N., Poduri, A., Chen, H.I., Goldstone, A.B., Woo, J., Fuller, G.G., Dunn, A.R. and Red-Horse, K. (2017). DACH1 stimulates shear stress-guided endothelial cell migration and coronary artery growth through the CXCL12–CXCR4 signaling axis. Genes & Development, 31(13), pp.1308–1324.

Chiang, I.K.-N., Fritzsche, M., Pichol-Thievend, C., Neal, A., Holmes, K., Lagendijk, A., Overman, J., D’Angelo, D., Omini, A., Hermkens, D., Lesieur, E., Fossat, N., Radziewic, T., Liu, K., Ratnayaka, I., Corada, M., Bou-Gharios, G., Tam, P.P.L., Carroll, J. and Dejana, E. (2017). Correction: SoxF factors induce Notch1 expression via direct transcriptional regulation during early arterial development. Development doi: 10.1242/dev.146241. Development, 144(20), pp.3847–3848.

Cho, C.-H., Lee, S.-Y., Shin, H.-S., Philipson, K.D. and Lee, C.O. (2003). Partial rescue of the Na+-Ca2+ exchanger (NCX1) knock-out mouse by transgenic expression of NCX1. Experimental & Molecular Medicine, 35(2), pp.125–135.

Chong, D.C., Koo, Y., Xu, K., Fu, S. and Cleaver, O. (2011). Stepwise arteriovenous fate acquisition during mammalian vasculogenesis. Developmental Dynamics, 240(9), pp.2153–2165.

Coan, P.M., Ferguson-Smith, A.C. and Burton, G.J. (2004). Developmental Dynamics of the Definitive Mouse Placenta Assessed by Stereology. Biology of Reproduction, 70(6), pp.1806–1813.

Courtney, J., Troja, W., Owens, K.J., Brockway, H.M., Hinton, A.C., Hinton, R.B., Cnota, J.F. and Jones, H.N. (2020). Abnormalities of placental development and function are associated with the different fetal growth patterns of hypoplastic left heart syndrome and transposition of the great arteries. Placenta, 101, pp.57–65.

Cross, J.C., Simmons, D.G. and Watson, E.D. (2003). Chorioallantoic Morphogenesis and Formation of the Placental Villous Tree. Annals of the New York Academy of Sciences, 995(1), pp.84–93.

Detmar, J., Rennie, M.Y., Whiteley, K.J., Qu, D., Taniuchi, Y., Shang, X., Casper, R.F., Adamson, S.L., Sled, J.G. and Jurisicova, A. (2008). Fetal growth restriction triggered by polycyclic aromatic hydrocarbons is associated with altered placental vasculature and AhR-dependent changes in cell death. American Journal of Physiology-Endocrinology and Metabolism, 295(2), pp.E519–E530.

Downs, K.M. and Rodriguez, A.M. (2019). The mouse fetal-placental arterial connection: A paradigm involving the primitive streak and visceral endoderm with implications for human development. WIREs Developmental Biology, 9(2).

Drake, C.J. and Fleming, P.A. (2000). Vasculogenesis in the day 6.5 to 9.5 mouse embryo. Blood, 95(5), pp.1671–1679.

Duarte, A. (2004). Dosage-sensitive requirement for mouse Dll4 in artery development. Genes & Development, 18(20), pp.2474–2478.

Freyer, L., Hsu, C.-W., Nowotschin, S., Pauli, A., Ishida, J., Kuba, K., Fukamizu, A., Schier, A.F., Hoodless, P.A., Dickinson, M.E. and Hadjantonakis, A.-K. (2017). Loss of Apela Peptide in Mice Causes Low Penetrance Embryonic Lethality and Defects in Early Mesodermal Derivatives. Cell Reports, 20(9), pp.2116–2130.

Gale, N.W., Dominguez, M.G., Noguera, I., Pan, L., Hughes, V., Valenzuela, D.M., Murphy, A.J., Adams, N.C., Lin, H.C., Holash, J., Thurston, G. and Yancopoulos, G.D. (2004). Haploinsufficiency of delta-like 4 ligand results in embryonic lethality due to major defects in arterial and vascular development. Proceedings of the National Academy of Sciences, 101(45), pp.15949–15954.

Gordon, E.J., Gale, N.W. and Harvey, N.L. (2008). Expression of the hyaluronan receptor LYVE-1 is not restricted to the lymphatic vasculature; LYVE-1 is also expressed on embryonic blood vessels. Developmental Dynamics: An Official Publication of the American Association of Anatomists, [online] 237(7), pp.1901–1909. Available at: https://pubmed.ncbi.nlm.nih.gov/18570254/ [Accessed 20 Mar. 2021].

Ho, L., Dijk, M. van, Chye, S.T.J., Messerschmidt, D.M., Chng, S.C., Ong, S., Yi, L.K., Boussata, S., Goh, G.H.-Y., Afink, G.B., Lim, C.Y., Dunn, N.R., Solter, D., Knowles, B.B. and Reversade, B. (2017). ELABELA deficiency promotes preeclampsia and cardiovascular malformations in mice. Science, [online] 357(6352), pp.707–713. Available at: https://science.sciencemag.org/content/357/6352/707/tab-article-info [Accessed 20 Mar. 2021].

Interactions between Trophoblast Cells and the Maternal and Fetal Circulation in the Mouse Placenta. (2002). Developmental Biology, [online] 250(2), pp.358–373. Available at: https://www.sciencedirect.com/science/article/pii/S0012160602907736 [Accessed 10 Mar. 2021].

Jones, H.N., Olbrych, S.K., Smith, K.L., Cnota, J.F., Habli, M., Ramos-Gonzales, O., Owens, K.J., Hinton, A.C., Polzin, W.J., Muglia, L.J. and Hinton, R.B. (2015). Hypoplastic left heart syndrome is associated with structural and vascular placental abnormalities and leptin dysregulation. Placenta, [online] 36(10), pp.1078–1086. Available at: https://pubmed.ncbi.nlm.nih.gov/26278057/ [Accessed 20 Mar. 2021].

Kalisch-Smith, J.I., Ved, N., Szumska, D., Munro, J., Troup, M., Harris, S.E., Jacquemot, A., Miller, J.J., Stuart, E.M., Wolna, M., Hardman, E., Prin, F., Lana-Elola, E., Aoidi, R., Fisher, E.M.C., Tybulewicz, V.L.J., Mohun, T.J., Lakhal-Littleton, S., Giannoulatou, E. and Sparrow, D.B. (2020). Maternal iron deficiency perturbs embryonic cardiovascular development. bioRXIV.

Kang, Y., Kim, J., Anderson, J.P., Wu, J., Gleim, S.R., Kundu, R.K., McLean, D.L., Kim, J., Park, H., Jin, S., Hwa, J., Quertermous, T. and Chun, H.J. (2013). Apelin-APJ Signaling Is a Critical Regulator of Endothelial MEF2 Activation in Cardiovascular Development. Circulation Research, 113(1), pp.22–31.

Lee, L., Ghorbanian, Y., Weissman, I., Inlay, M. and Mikkola, H. (2015). Lyve1 marks yolk sac definitive hemogenic endothelium. Experimental Hematology, 43(9), p.S80.

Levin, H.I., Sullivan-Pyke, C.S., Papaioannou, V.E., Wapner, R.J., Kitajewski, J.K., Shawber, C.J. and Douglas, N.C. (2017). Dynamic maternal and fetal Notch activity and expression in placentation. Placenta, 55, pp.5–12.

Lupu, I.-E., Redpath, A.N. and Smart, N. (2020). Spatiotemporal Analysis Reveals Overlap of Key Proepicardial Markers in the Developing Murine Heart. Stem Cell Reports, 14(5), pp.770–787.

Madisen, L., Zwingman, T.A., Sunkin, S.M., Oh, S.W., Zariwala, H.A., Gu, H., Ng, L.L., Palmiter, R.D., Hawrylycz, M.J., Jones, A.R., Lein, E.S. and Zeng, H. (2009). A robust and high-throughput Cre reporting and characterization system for the whole mouse brain. Nature Neuroscience, 13(1), pp.133–140.

Marsh, B. and Blelloch, R. (2020). Single nuclei RNA-seq of mouse placental labyrinth development. [online] eLife. Available at: https://elifesciences.org/articles/60266v1 [Accessed 20 Mar. 2021].

Maslen, C.L. (2018). Recent Advances in Placenta–Heart Interactions. Frontiers in Physiology, 9.

Matthiesen, N.B., Henriksen, T.B., Agergaard, P., Gaynor, J.W., Bach, C.C., Hjortdal, V.E. and Østergaard, J.R. (2016). Congenital Heart Defects and Indices of Placental and Fetal Growth in a Nationwide Study of 924 422 Liveborn Infants. Circulation, 134(20), pp.1546– 1556.

Moreau, J.L.M., Artap, S.T., Shi, H., Chapman, G., Leone, G., Sparrow, D.B. and Dunwoodie, S.L. (2014). Cited2 is required in trophoblasts for correct placental capillary patterning. Developmental Biology, 392(1), pp.62–79.

Natale, B.V., Mehta, P., Vu, P., Schweitzer, C., Gustin, K., Kotadia, R. and Natale, D.R.C. (2018). Reduced Uteroplacental Perfusion Pressure (RUPP) causes altered trophoblast differentiation and pericyte reduction in the mouse placenta labyrinth. Scientific Reports, 8(1).

Navankasattusas, S., et al. “The Netrin Receptor UNC5B Promotes Angiogenesis in Specific Vascular Beds.” Development, vol. 135, no. 4, 15 Feb. 2008, pp. 659–667, 10.1242/dev.013623. Accessed 29 Mar. 2021.

Nelson, A.C., Mould, A.W., Bikoff, E.K. and Robertson, E.J. (2016). Single-cell RNA-seq reveals cell type-specific transcriptional signatures at the maternal–foetal interface during pregnancy. Nature Communications, 7, p.11414.

Outhwaite, J.E., Patel, J. and Simmons, D.G. (2019). Secondary Placental Defects in Cxadr Mutant Mice. Frontiers in Physiology, 10.

Perez-Garcia, V., Fineberg, E., Wilson, R., Murray, A., Mazzeo, C.I., Tudor, C., Sienerth, A., White, J.K., Tuck, E., Ryder, E.J., Gleeson, D., Siragher, E., Wardle-Jones, H., Staudt, N., Wali, N., Collins, J., Geyer, S., Busch-Nentwich, E.M., Galli, A. and Smith, J.C. (2018). Placentation defects are highly prevalent in embryonic lethal mouse mutants. Nature, 555(7697), pp.463–468.

Pham, T.H.M., Baluk, P., Xu, Y., Grigorova, I., Bankovich, A.J., Pappu, R., Coughlin, S.R., McDonald, D.M., Schwab, S.R. and Cyster, J.G. (2009). Lymphatic endothelial cell sphingosine kinase activity is required for lymphocyte egress and lymphatic patterning. Journal of Experimental Medicine, [online] 207(1), pp.17–27. Available at: https://rupress.org/jem/article/207/1/17/40557/Lymphatic-endothelial-cell-sphingosine-kinase [Accessed 20 Mar. 2021].

Raftrey, B., Williams, I., Rios Coronado, P.E., Chang, A.H., Zhao, M., Roth, R., Racelis, R., D’Amato, G., Phansalkar, R., Gonzalez, K.M., Zhang, Y., Bernstein, D. and Red-Horse, K. (2020). Dach1 extends artery networks and protects against cardiac injury. bioRvix. [online] Available at: https://doi.org/10.1101/2020.08.07.242164 [Accessed 10 Mar. 2021].

Rennie, M.Y., Cahill, L.S., Adamson, S.L. and Sled, J.G. (2017). Arterio-venous fetoplacental vascular geometry and hemodynamics in the mouse placenta. Placenta, [online] 58, pp.46–51. Available at: https://doi.org/10.1016/j.placenta.2017.08.007 [Accessed 10 Mar. 2021].

Rychik, J., Goff, D., McKay, E., Mott, A., Tian, Z., Licht, D.J. and Gaynor, J.W. (2018). Characterization of the Placenta in the Newborn with Congenital Heart Disease: Distinctions Based on Type of Cardiac Malformation. Pediatric Cardiology, 39(6), pp.1165– 1171.

Sacilotto, N., Chouliaras, K.M., Nikitenko, L.L., Lu, Y.W., Fritzsche, M., Wallace, M.D., Nornes, S., García-Moreno, F., Payne, S., Bridges, E., Liu, K., Biggs, D., Ratnayaka, I., Herbert, S.P., Molnár, Z., Harris, A.L., Davies, B., Bond, G.L., Bou-Gharios, G. and Schwarz, J.J. (2016). MEF2 transcription factors are key regulators of sprouting angiogenesis. Genes & Development, 30(20), pp.2297–2309.

Sacilotto, N., Monteiro, R., Fritzsche, M., Becker, P.W., Sanchez-del-Campo, L., Liu, K., Pinheiro, P., Ratnayaka, I., Davies, B., Goding, C.R., Patient, R., Bou-Gharios, G. and De Val, S. (2013). Analysis of Dll4 regulation reveals a combinatorial role for Sox and Notch in arterial development. Proceedings of the National Academy of Sciences, 110(29), pp.11893–11898.

Saint-Geniez, M., Masri, B., Malecaze, F., Knibiehler, B. and Audigier, Y. (2002). Expression of the murine msr/apj receptor and its ligand apelin is upregulated during formation of the retinal vessels. Mechanisms of Development, 110(1-2), pp.183–186.

Sharma, B., Ho, L., Ford, G.H., Chen, H.I., Goldstone, A.B., Woo, Y.J., Quertermous, T., Reversade, B. and Red-Horse, K. (2017). Alternative Progenitor Cells Compensate to Rebuild the Coronary Vasculature in Elabela-and Apj-Deficient Hearts. Developmental Cell, 42(6), pp.655–666.e3.

Shaut, C.A.E., Keene, D.R., Sorensen, L.K., Li, D.Y. and Stadler, H.S. (2008). HOXA13 Is Essential for Placental Vascular Patterning and Labyrinth Endothelial Specification. PLoS Genetics, 4(5), p.e1000073.

Shi, H., O’Reilly, V.C., Moreau, J.L.M., Bewes, T.R., Yam, M.X., Chapman, B.E., Grieve, S.M., Stocker, R., Graham, R.M., Chapman, G., Sparrow, D.B. and Dunwoodie, S.L. (2016). Gestational stress induces the unfolded protein response, resulting in heart defects. Development, 143(14), pp.2561–2572.

Simmons, D.G., Natale, D.R.C., Begay, V., Hughes, M., Leutz, A. and Cross, J.C. (2008). Early patterning of the chorion leads to the trilaminar trophoblast cell structure in the placental labyrinth. Development, [online] 135(12), pp.2083–2091. Available at: https://dev.biologists.org/content/135/12/2083.short [Accessed 10 Mar. 2021].

Tai-Nagara, I., Yoshikawa, Y., Numata, N., Ando, T., Okabe, K., Sugiura, Y., Ieda, M., Takakura, N., Nakagawa, O., Zhou, B., Okabayashi, K., Suematsu, M., Kitagawa, Y., Bastmeyer, M., Sato, K., Klein, R., Navankasattusas, S., Li, D.Y., Yamagishi, S. and Kubota, Y. (2017). Placental labyrinth formation in mice requires endothelial FLRT2/UNC5B signaling. Development, 144(13), pp.2392–2401.

Thiery, J.P. (2002). Epithelial–mesenchymal transitions in tumour progression. Nature Reviews Cancer, 2(6), pp.442–454.

Thornburg, K.L., O’Tierney, P.F. and Louey, S. (2010). Review: The Placenta is a Programming Agent for Cardiovascular Disease. Placenta, 31, pp.S54–S59.

Wang, H.U., Chen, Z.-F. and Anderson, D.J. (1998). Molecular Distinction and Angiogenic Interaction between Embryonic Arteries and Veins Revealed by ephrin-B2 and Its Receptor Eph-B4. Cell, 93(5), pp.741–753.

Wang, Y., Nakayama, M., Pitulescu, M.E., Schmidt, T.S., Bochenek, M.L., Sakakibara, A., Adams, S., Davy, A., Deutsch, U., Lüthi, U., Barberis, A., Benjamin, L.E., Mäkinen, T., Nobes, C.D. and Adams, R.H. (2010). Ephrin-B2 controls VEGF-induced angiogenesis and lymphangiogenesis. Nature, 465(7297), pp.483–486.

WHO. (2015). WHO | The global prevalence of anaemia in 2011. [online] Available at: https://www.who.int/nutrition/publications/micronutrients/global_prevalence_anaemia_2011/en/.

Withington, S.L., Scott, A.N., Saunders, D.N., Lopes Floro, K., Preis, J.I., Michalicek, J., Maclean, K., Sparrow, D.B., Barbera, J.P.M. and Dunwoodie, S.L. (2006). Loss of Cited2 affects trophoblast formation and vascularization of the mouse placenta. Developmental Biology, 294(1), pp.67–82.

Wyrwoll, C.S., Noble, J., Thomson, A., Tesic, D., Miller, M.R., Rog-Zielinska, E.A., Moran, C.M., Seckl, J.R., Chapman, K.E. and Holmes, M.C. (2016). Pravastatin ameliorates placental vascular defects, fetal growth, and cardiac function in a model of glucocorticoid excess. Proceedings of the National Academy of Sciences, [online] 113(22), pp.6265– 6270. Available at: https://www.pnas.org/content/113/22/6265 [Accessed 20 Mar. 2021].

Wythe, Joshua D., Dang, Lan T.H., Devine, W. Patrick, Boudreau, E., Artap, Stanley T., He, D., Schachterle, W., Stainier, Didier Y.R., Oettgen, P., Black, Brian L., Bruneau, Benoit G. and Fish, Jason E. (2013). ETS Factors Regulate Vegf-Dependent Arterial Specification. Developmental Cell, 26(1), pp.45–58.

Zudaire, E., Gambardella, L., Kurcz, C. and Vermeren, S. (2011). A Computational Tool for Quantitative Analysis of Vascular Networks. PLoS ONE, 6(11), p.e27385.

